# Comprehensive analysis of microsatellite polymorphisms in human populations

**DOI:** 10.1101/2022.06.08.495243

**Authors:** Leo Gochi, Yosuke Kawai, Akihiro Fujimoto

## Abstract

Microsatellites (MS) are tandem repeats of short units and have been used for population genetics, individual identification, and medical genetics. However, studies of MS on a whole genome level are limited, and genotyping methods for MS have yet to be established. Here, we analyzed approximately 8.5 million MS regions using a previously developed MS caller (MIVcall method) for three large publicly available human genome sequencing data sets: the Korean Personal Genome Project (KPGP), Simons Genome Diversity Project (SGDP), and Human Genome Diversity Project (HGDP). Our analysis identified 253,114 polymorphic MS. A comparison among different populations suggests that MS in the coding region evolved by random genetic drift and natural selection. In an analysis of genetic structures, MS clearly revealed population structures as SNPs and detected clusters that were not found by SNPs in African and Oceanian populations. Based on the MS polymorphisms, we selected an effective MS set for individual identification. We also showed that our MS analysis method can be applied to ancient DNA samples. This study provides a comprehensive picture of MS polymorphisms and application to human population studies.

## Introduction

Repetitive sequences account for more than two-thirds of the human genome (1). Among them, sequences consisting of tandem repeats of short units are classified as microsatellites (MS). The mutation rate of MS is higher than these of other genomic regions, and MS have high diversity among individuals (1).

Due to their high heterozygosity and multiallelic nature, MS have been widely used as genetic markers in population studies (2–5). Previous studies analyzed dozens to several hundreds of MS and revealed the genetic structure of modern human populations, the relationship of genetic and linguistic variations, and the trace of natural selection (3–6). These studies have made a great contribution to understanding human evolutionary history. However, conventional PCR-based MS genotyping methods are not suitable for high-throughput genotyping; therefore, MS has largely been replaced with single nucleotide polymorphism (SNP) in related studies. Indeed, in the past decade, genome-wide SNP analysis had become a standard method for human population genetics (7,8).

In addition to population studies, MS has been widely used for personal identification in forensic science and paternity testing (9,10) since MS are multiallelic and have more discriminative power than SNP (2). Established common MS sets, such as the Globalfiler kit, have been used for many years (11). Although these MS marker sets have sufficient power for most cases, they are not selected from entire genomes and there may be other more efficient MS in the human genome.

Next generation sequencing technologies (NGS) enable whole genome sequencing (WGS). In the past decade, applications of NGS and the development of algorithms for the analysis have successfully identified genetic variations, including single nucleotide variations, insertions and deletions, copy number variations, and structural variations (12). However, due to the short read length and the high sequencing error rate in repeat regions, the identification of mutations and polymorphisms in MS regions have been difficult. Previously, several groups including ours have developed MS genotyping tools from WGS data (13–15). These methods identified somatic mutations or germline polymorphisms in MS in the human population and revealed genome-wide patterns of MS polymorphisms, factors that determine the mutation rate of MS, and functional roles of MS on gene expressions (13–17). Although these studies provide important information on MS, the MS genotyping method is not perfect, and only few studies have been conducted for MS polymorphisms. Indeed, the patterns of genome-wide MS polymorphisms have not been well analyzed in various human populations. Nor has the amount of genetic variation in MS among different human populations been compared in detail. Furthermore, clustering based on principal component analysis (PCA) with MS polymorphisms has presented unclear results compared to that with SNPs (14,18), and the efficiency of genome-wide MS for analyzing population structures requires more study.

Here, we analyzed approximately nine million MS regions using a previously developed MS caller for three large publicly available human genome sequencing data sets: Korean Personal Genome Project (KPGP), Simons Genome Diversity Project (SGDP), and Human Genome Diversity Project (HGDP) (13,18,19). We revealed the pattern of MS polymorphisms, analyzed the genetic structure of the populations with several dimensionality reduction methods, and identified useful candidate MS for individual identification. Additionally, we analyzed MS of an ancient DNA sample (20). Our analysis provides a comprehensive picture of MS polymorphisms and their application to human population studies.

## Results

### Establishment of MS calling

We identified genotypes of MS with the MIVcall method (13). MIVcall outputs the log10(likelihood) and number of reads for each MS locus, which can be used to evaluate the reliability of MS genotypes. We examined these parameters using monozygotic twins in the KPGP (KPGP-00088 and KPGP-00089). Because monozygotic twins have completely same genotypes, we considered all disconcordant genotypes between the twins as errors. We compared the genotypes of 8,343,174 MS in the twins and classified them into concordant homozygote, concordant heterozygote, disconcordance of two alleles, and disconcordance of one allele (S1 Table). Based on the result, the cutoff -log10(likelihood) and minimum number of reads for allele detection were set to -4 and 3, respectively, in this study (S1 Table).

### Selection of samples and MS

We selected samples for the analysis. In this study, MS covered by less than 10 reads were considered insufficient depth, and we excluded samples if more than 4% had insufficient depth of MS. As a result, 277 samples from the SGDP, 692 samples from the HGDP, and 81 samples from the KPGP (excluding one of the monozygotic twins) were selected.

We next selected MS loci for this study using the 81 Korean samples from the KPGP. We selected MS that were genotyped (number of reads ≥ 10) in more than 85% of the 81 samples, leaving 8,468,218 MS. We also tested deviation from the Hardy-Weinberg equilibrium (HWE) in the KPGP, but no MS showed significant deviation.

Of the selected 8,468,218 MS, 7,740,569 MS were monomorphic. For each MS, minor allele frequency (MAF) was calculated by 1 – (major allele frequency). Of these, 727,649 MS had variations in at least one sample, and 253,114 had a MAF ≥ 1% (S1 Fig). Of all MS, 71,040 MS were in coding sequences (CDS), and 8,397,178 MS were in non-CDS (S2 Table). Among the CDS MS, 893 MS had variations in at least one sample, and 71 MS had a MAF ≥ 1% (S2 Table).

### Genome-wide pattern of MS polymorphisms

The length of MS was negatively correlated to the number of samples with insufficient depth (p-value < 10^−200^ Kruskal-Wallis test) but positively correlated to the number of alleles (p-value < 10^−200^ Kruskal-Wallis test) (Fig 1A and B). The increase in the number of alleles was more gradual in longer MS (Fig 1B). The longer MS had fewer reads fully covering them compared to shorter MS and thus a lower detection sensitivity. The number and heterozygosity of different repeat units showed that MS with higher AT content had significantly higher heterozygosity (Pearson’s uncorrelated test p-value = 1.47×10^−61^), suggesting that the mutability of MS is affected by the base composition (Fig 1 C and D).

**Fig. 1.**
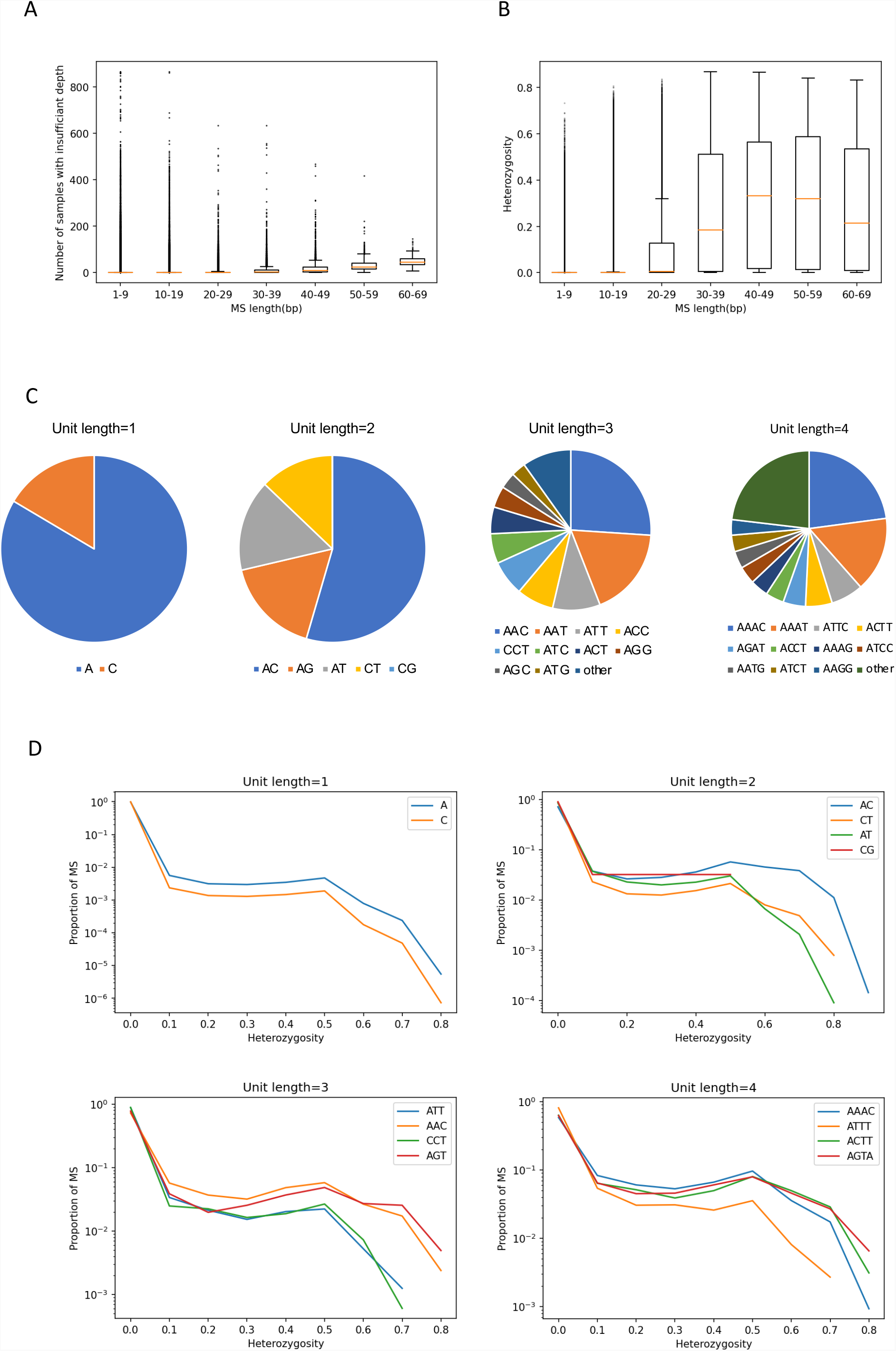
Features of microsatellites (MS) in the human genome. (A) MS length and number of samples with insufficient depth. The MS length was negatively correlated to the number of samples with insufficient depth (p-value < 10^−200^ Kruskal-Wallis test). (B) MS length and heterozygosity. MS length was positively correlated to the number of alleles (p-value < 10^−200^ Kruskal-Wallis test). (C) Proportion of MS unit types. (D) Number of MS and heterozygosity of different units. MS with higher AT content had significantly higher heterozygosity (Pearson’s uncorrelated test p-value = 1.47×10^−61^).

A comparison between CDS and non-CDS showed that MS in CDS had a higher GC content (p-value = 6.49×10^−144^ Wilcoxon rank sum test), shorter length (p-value = 1.25×10^−29^ Wilcoxon rank sum test), smaller number of alleles (p-value = 1.80×10^−47^ Wilcoxon rank sum test), and lower heterozygosity (p-value = 1.42×10^−89^ Wilcoxon rank sum test) (Fig 2 A-D). A comparison of distributions of the number MS in each unit length showed that polymorphic 3*n* MS (3 and 6 bp) were frequent in the CDS region (p-value = 5.22×10^−232^ Fisher’s exact test) (Fig 2 E-H, S3 Table). In the CDS, 3*n* MS had higher heterozygosity than non-3*n* MS (1, 2, 4, 5 and 7 bp) (p-value = 5.23×10^−13^ Wilcoxon rank sum test), suggesting that non-3*n* MS was strongly influenced by negative selection (Fig 2I).

**Fig. 2.**
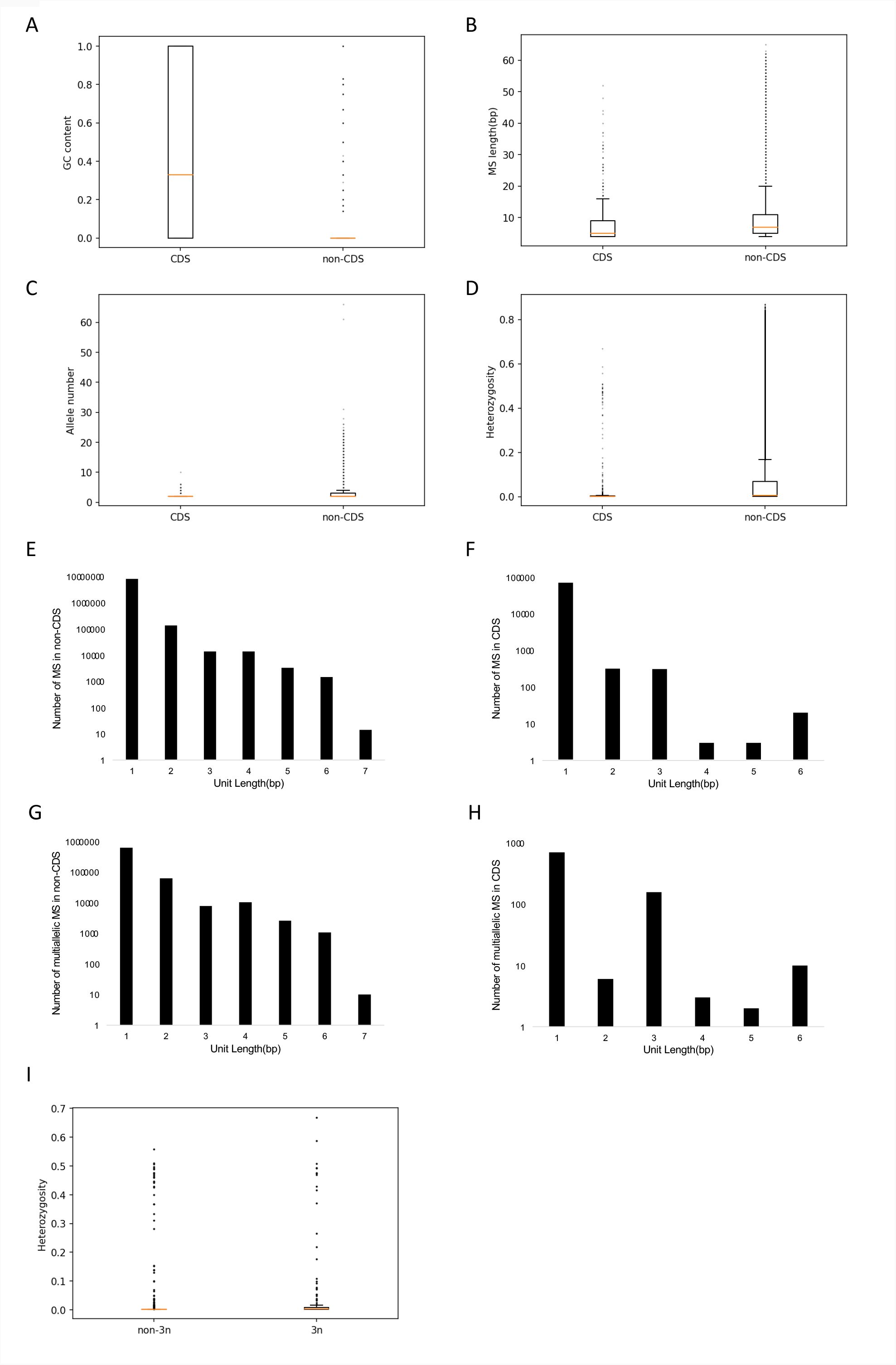
Features of MS in CDS and non-CDS regions. (A) MS in CDS had higher GC content (p-value = 6.49×10^−144^ Wilcoxon rank sum test). (B) MS in CDS had shorter length (p-value = 1.25×10^−29^ Wilcoxon rank sum test). (C) MS in CDS had fewer alleles (p-value = 1.80×10^−47^ Wilcoxon rank sum test). (D) MS in CDS had lower heterozygosity (p-value = 1.42×10^−89^ Wilcoxon rank sum test). (E) Total number of MS loci per unit length in non-CDS. (F) Total number of MS loci per unit length in CDS. (G) Number of multiallelic MS of each unit length in CDS. (H) Number of multiallelic MS of each unit length in non-CDS. (I) Heterozygosity of MS in non-CDS and CDS. Non-3*n* MS had lower heterozygosity than 3*n* MS (p-value = 5.23×10^−13^ Wilcoxon rank sum test).

To compare the genetic variation among human populations, we calculated the distribution of the heterozygosity of all MS, MS in CDS, MS in non-CDS, 3*n* MS in CDS, and non-3*n* MS in CDS (Fig 3 A-E and S4 Table). We also calculated the ratio of the mean heterozygosity of non-3*n* MS to mean heterozygosity of 3*n* MS in CDS among each region (Fig 3F). Africa had the highest heterozygosity and lowest ratio of mean heterozygosity (Fig 3 and S4 Table).

**Fig. 3.**
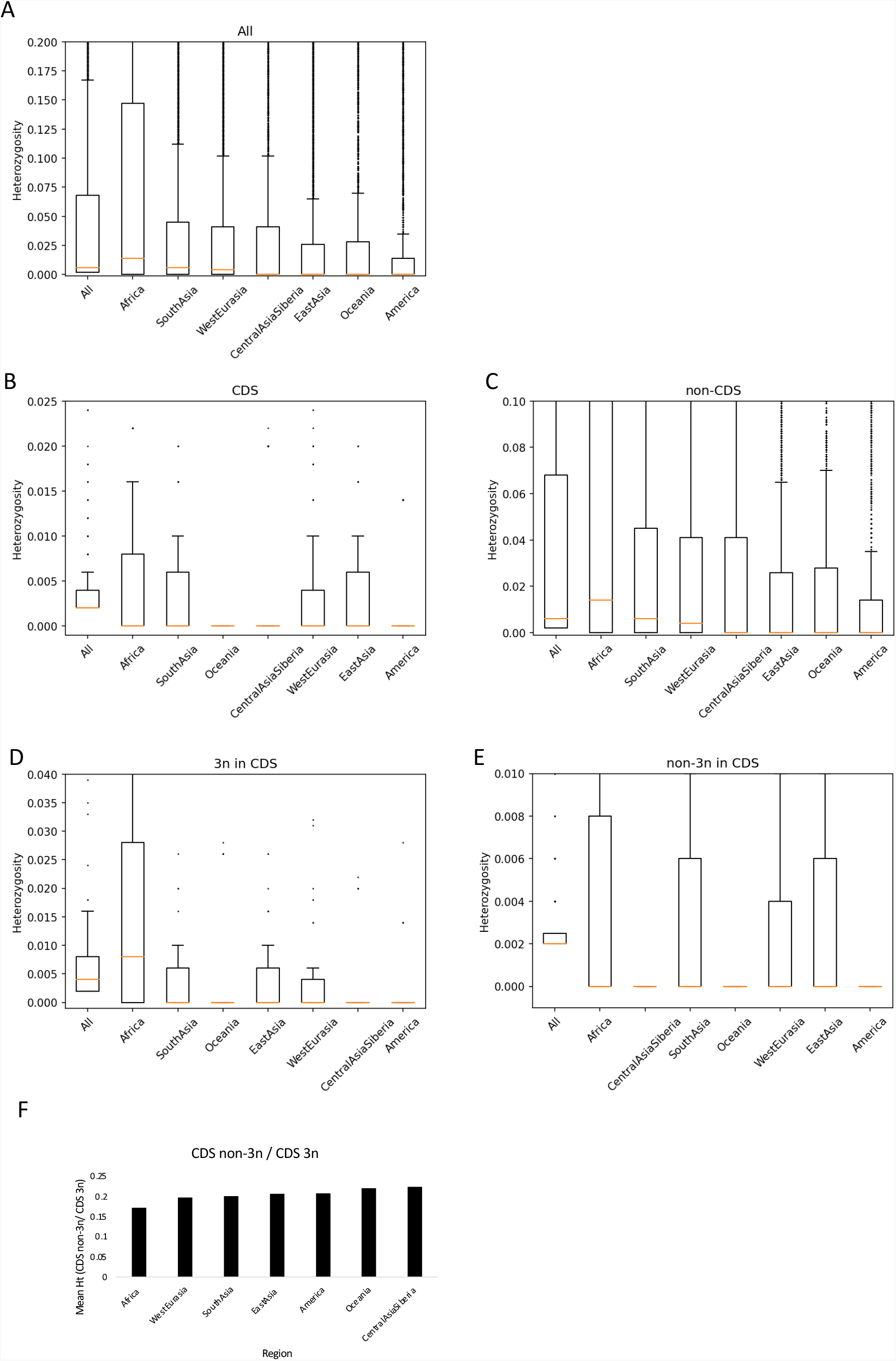
Heterozygosity of MS among demographic regions. (A) Heterozygosity of all MS. (B) Heterozygosity of MS in CDS. (C) Heterozygosity of MS in non-CDS. (D) Heterozygosity of 3*n* MS in CDS. (E) Heterozygosity of non-3*n* MS in CDS. In (A)-(E), regions were sorted by their mean heterozygosity. (F) Ratios of mean heterozygosity of non-3*n* MS to 3*n* MS in CDS.

We observed that 689 genes had 724 polymorphic non-3*n* MS (S4 Table, S5 Table). We performed a pathway analysis for these genes but did not find any significantly over-represented pathway (data not shown).

### Analysis of population structure

To analyze the population structure, we conducted five dimensionality reduction methods for MS polymorphisms: PCA, t-Distributed Stochastic Neighbor Embedding (t-SNE), Uniform Manifold Approximation and Projection (UMAP), PCA-t-SNE, and PCA-UMAP using MS and SNP. For this analysis, we used MS and SNP with MAF ≥ 1%. In the MS, 253,114 MS had MAF ≥ 1% in all samples, 340,114 in Africa, 176,214 in America, 229,636 in Central Asia and Siberia, 197,081 in East Asia, 231,706 in Oceania, 220,737 in South Asia, and 209,204 in West Eurasia (Supplementary data1). In the SNPs, 486,579 SNPs had MAF ≥ 1% in all samples, 515,456 in Africa, 358,538 in America, 416,191 in Central Asia and Siberia, 403,015 in East Asia, 389,358 in Oceania, 437,410 in South Asia, and 432,163 in West Eurasia (S2 Table).

To apply dimensionality reduction methods, we converted MS genotypes to numerical values using two methods, multiallelic method and average method (see Methods), and performed PCA with both methods for all samples. Although the results were not significantly different between the two, the average method had a higher contribution rate (PC1 = 5.80%) than the multiallelic method (PC1 = 4.97%) (S2 Fig). Therefore, we selected the average method in this study.

PCA was conducted for all regions and each individual region (Fig 4, S3 Fig). We found similar patterns between MS and SNPs for all samples (Fig 4 AB). However, patterns were slightly different in African and Oceanian populations between MS and SNPs (Fig 4 C-F). These results suggest that genome-wide MS has compatible resolution to SNPs for the genetic structure of human populations and that MS can be used to find new genetic structures. t-SNE, UMAP, PCA-t-SNE, and PCA-UMAP were also conducted for all samples (S4 Fig). In most of these analyses MS did not detect novel clusters; however, MS discriminated African populations from the others in the t-SNE analysis (S4 Fig A).

**Fig. 4.**
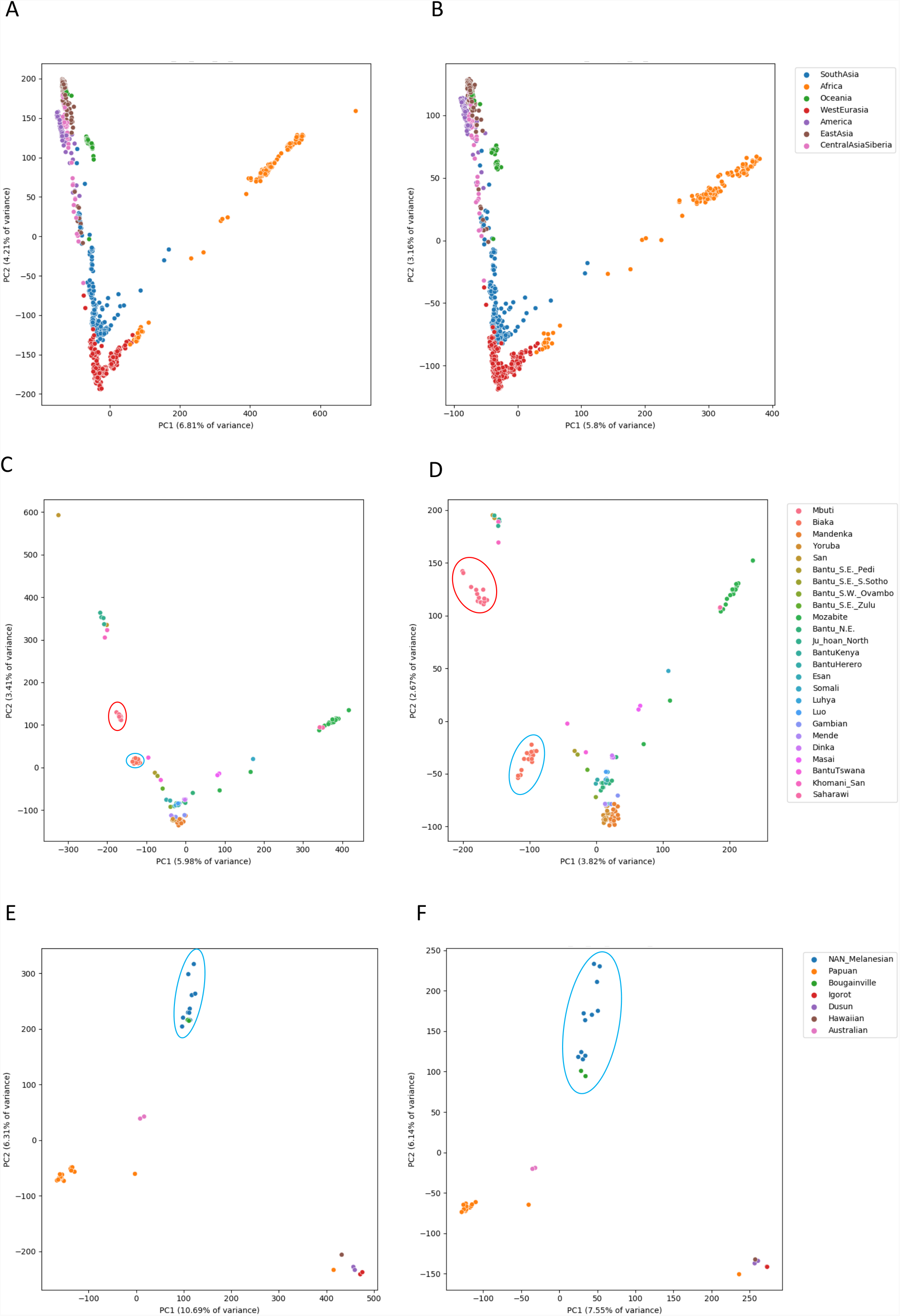
PCA using MS and SNPs. (A) PCA for all samples using MS. (B) PCA for all samples using SNPs. (C) PCA for African populations using MS. (D) PCA for African populations using SNPs. PC2 values of the Mbuti and Biaka populations were different between MS and SNP. (E) PCA for Oceanian populations using MS. (F) PCA for Oceanian populations using SNPs. NAN_Melanesuans and Bougainville were clustered together in the SNP but separated in the MS.

### MS marker set for individual identification

We selected a set of 22 MS and calculated the discriminative power. From MS with 4-bp units, 22 MS with the highest heterozygosity were selected from each chromosome (Table 1).

**Table 1.** MS set for individual identification

| Chromosome | Start position | End position | Type | Max length of MS<br>in ICGC Japanese<br>samples (bp) | Max length of MS in<br>HGDP+SGDP (bp) | Number of alleles in<br>ICGC Japanese<br>samples | Number of alleles<br>in HGDP+SGDP | Match probability<br>in ICGC Japanese<br>samples | Match<br>probability in<br>HGDP+SGDP |
| --- | --- | --- | --- | --- | --- | --- | --- | --- | --- |
| 1 | 105083241 | 105083272 | (TATC)n | 48 | 55 | 12 | 20 | 0.074 | 0.076 |
| 2 | 11032690 | 11032717 | (CTTC)n | 44 | 56 | 5 | 10 | 0.270 | 0.134 |
| 3 | 76233694 | 76233746 | (AGAT)n | 61 | 62 | 11 | 19 | 0.075 | 0.107 |
| 4 | 117885601 | 117885630 | (TATC)n | 49 | 53 | 11 | 16 | 0.085 | 0.054 |
| 5 | 82900377 | 82900424 | (TCTA)n | 55 | 55 | 10 | 14 | 0.154 | 0.095 |
| 6 | 49948157 | 49948210 | (ATCT)n | 66 | 70 | 10 | 11 | 0.142 | 0.091 |
| 7 | 67348144 | 67348171 | (TTTC)n | 44 | 60 | 7 | 17 | 0.318 | 0.150 |
| 8 | 108769208 | 108769243 | (TAAG)n | 56 | 60 | 7 | 13 | 0.247 | 0.165 |
| 9 | 101627740 | 101627784 | (ATCT)n | 57 | 61 | 7 | 8 | 0.231 | 0.178 |
| 10 | 2918258 | 2918301 | (TAGA)n | 52 | 48 | 7 | 9 | 0.175 | 0.095 |
| 11 | 114544813 | 114544850 | (AAAC)n | 47 | 47 | 10 | 14 | 0.209 | 0.149 |
| 12 | 121657632 | 121657667 | (TAGA)n | 52 | 52 | 9 | 11 | 0.189 | 0.116 |
| 13 | 107091029 | 107091073 | (ATGG)n | 50 | 54 | 5 | 9 | 0.187 | 0.148 |
| 14 | 38581137 | 38581184 | (ATAG)n | 58 | 58 | 12 | 15 | 0.171 | 0.122 |
| 15 | 47067031 | 47067072 | (GAAT)n | 50 | 46 | 7 | 12 | 0.202 | 0.143 |
| 16 | 86386308 | 86386351 | (GATA)n | 56 | 60 | 7 | 9 | 0.156 | 0.155 |
| 17 | 72561544 | 72561595 | (CTAT)n | 56 | 56 | 5 | 10 | 0.204 | 0.155 |
| 18 | 5249011 | 5249068 | (AGAT)n | 70 | 74 | 12 | 21 | 0.180 | 0.103 |
| 19 | 15754216 | 15754267 | (ATAG)n | 60 | 60 | 8 | 15 | 0.164 | 0.116 |
| 20 | 5596482 | 5596528 | (TATG)n | 47 | 51 | 7 | 9 | 0.178 | 0.126 |
| 21 | 42031280 | 42031319 | (TATC)n | 52 | 52 | 9 | 13 | 0.136 | 0.157 |
| 22 | 36513967 | 36514013 | (TATC)n | 57 | 75 | 13 | 31 | 0.194 | 0.114 |
| | | | | | | | Product of the<br>match probability<br>of all MS | $1.0 \times 10^{-17}$ | $6.3 \times 10^{-21}$ |

For the selected 22 loci, we calculated the discriminative power using allele frequencies of 255 Japanese ICGC datasets (Table 1). The discriminative power for this MS set was estimated to be 1.0×10^−17^.

### MS analysis for an ancient DNA sample

Although the genome-wide MS analysis of ancient DNA samples has yet to be conducted, an analysis of MS variation in ancient DNA samples may contribute to clarifying the genetic structure of ancient populations. Since low-quality sequencing data of ancient DNA samples can result in incorrect results, we selected an ancient DNA sample with high sequence depth (sample ID; F23) (20) and analyzed the distribution of the variant allele frequency (VAF) for MS (Fig 5 A-D). VAF was calculated by (number of reads with second most frequent pattern)/(total number of reads) for heterozygous MS. The distributions of VAF were quite different between F23 and modern human samples in MS with lengths < 3 bp, suggesting that genotypes contained many errors. However, in MS with lengths ≥ 3 bp, the distributions of VAF were not different (Fig 5 A-D). Therefore, we used MS with lengths ≥ 3 bp and performed PCA with HGDP and SGDP samples (All, East Asia, America, Oceania, Central Asia and Siberia, and South Asia) (Fig 5 E,F). F23 was clustered close to Central Asia and Siberia and East Asia populations in the PCA plot.

**Fig. 5.**
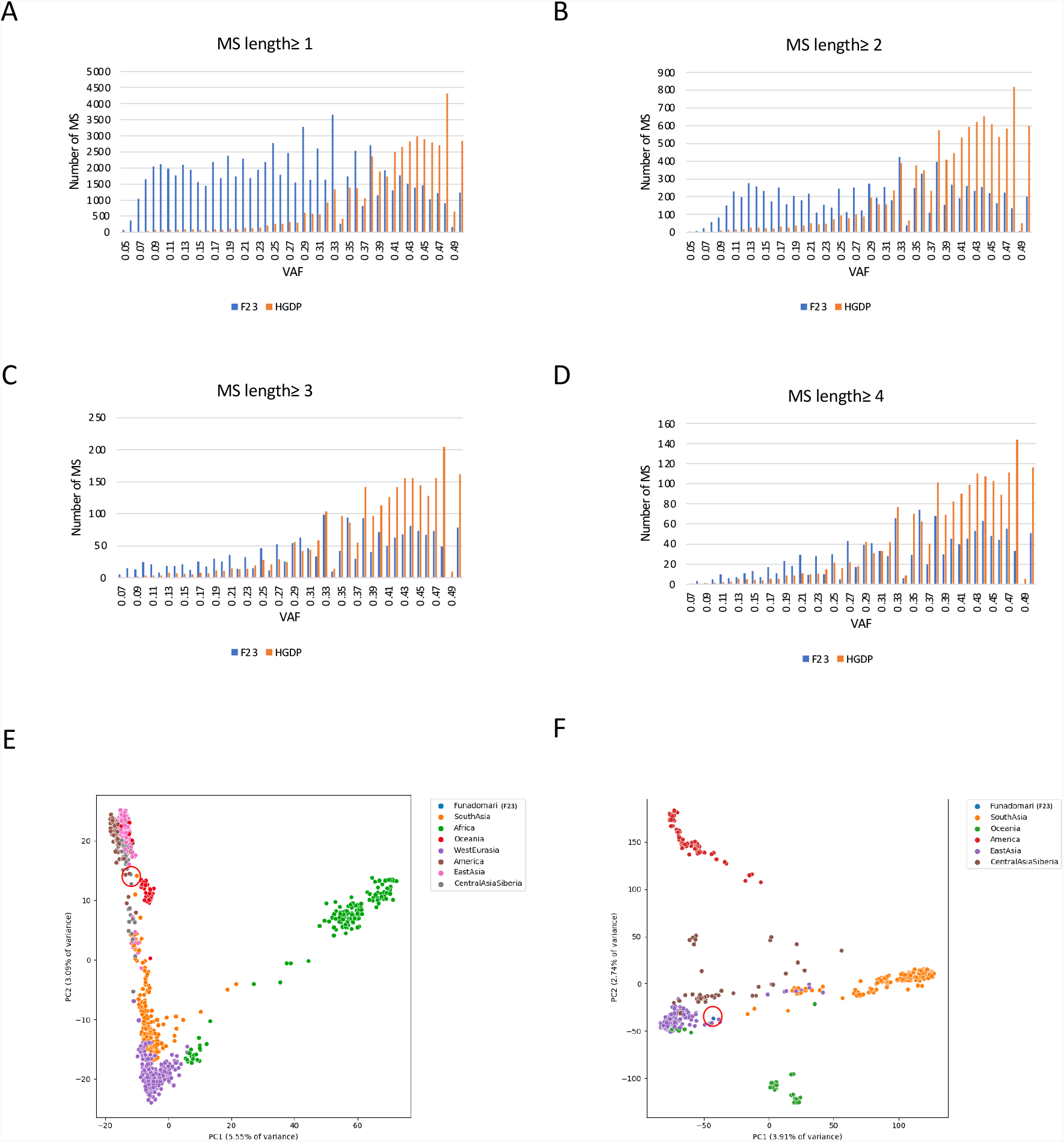
Analysis of an ancient sample (F23). Distribution of VAF in F23 and HGDP samples for unit lengths ≥ 1 bp (A), ≥ 2 bp (B), ≥ 3 bp (C), and ≥ 4 bp (D). The distributions of VAF were quite different between F23 and modern human samples in MS with lengths ≤ 2 bp. (E) PCA using MS for F23 and all modern human samples. (F) PCA using MS for F23 and modern human samples in South Asia, Oceania, America, East Asia, and Central Asia and Siberia.

## Discussion

Population genetic studies strongly depend on variant calling. Thus, like other types of variants, MS analysis is affected by genotyping methods. Since most MS calling methods analyze only predefined MS regions, the numbers of target MS are different among studies (700,000 loci in Gymrek et al., 2017 and Willems et al., 2014, and 1.6 million loci in Jakubosky, Smith, et al., 2020) (15,18). In this study, we analyzed a larger number of MS regions (8,468,218 MS) than previous studies for a comprehensive analysis of human MS polymorphisms. Compared to conventional PCR-based MS studies(17), this study has another advantage: MS were not influenced by ascertainment bias. Most conventional studies have analyzed pre-screened MS marker sets (21), which are influenced by the MS marker selection. On the other hand, we did not select MS based on the allele frequency in certain populations and could analyze features of MS and compare the variation among different populations without the influence of ascertainment bias.

We first selected parameters for the analysis based on the evaluation of monozygotic twins in the KPGP. The concordant rate of this parameter set was estimated to be 99.87%, which is sufficiently accurate for population studies (S1 Table). We then selected 8,468,218 MS based on the call rate and HWE in the KPGP for 81 Korean individuals. Our MS calling identified 253,114 MS with MAF ≥ 1% in the SGDP and HGDP (S1 Fig). Since the SGDP and HGDP datasets represent human genome diversity, these MS polymorphisms can be used for future population studies (S1 data). Previous genome-wide studies reported that MS polymorphisms could influence gene expression patterns and the risk of human diseases (17,22). Therefore applications of our MS set and our MS calling method may contribute to discovering novel disease susceptibility genes. In particular, 696 genes with non-3*n* MS are good targets for disease studies (S5 Table).

The amount of genetic variation reflects the effective population size (*N*). A comparison of heterozygosities across regions showed Africa with the highest (Fig 3, S4 Table), and America, which was estimated to have a very small effective population size, showed the lowest (Fig 3, S4 Table) (23). Such patterns were observed in other genetic variations and MS in a previous study (14,24), suggesting that the heterozygosity of MS reflects the size of each population. Theoretical population genetics also predicts that the effectiveness of natural selection depends on the selection coefficient (*s*) of the genetic variation and population size (25). Therefore, a comparison of genetic variations of MS in the CDS may provide additional information about the evolution of MS. Most of the 3*n* MS should not cause severe damage to protein functions and have neutral or nearly neutral effects, whereas non-3*n* MS cause a frameshift and should have deleterious effects. We attempted to evaluate the strength of negative selection among populations. For this purpose, we compared the average heterozygosity of non-3*n* MS in CDS with the average heterozygosity of 3*n* MS in CDS among populations (Fig 3F, S4 Table). The African population showed the lowest ratio, and populations with lower heterozygosity tended to have a higher ratio (Fig 3F, S4 Table). This pattern indicates that the African population has the largest effective population size and that stronger natural selection has acted to remove deleterious non-3*n* MS. In Central Asia and Siberia, the heterozygosity was not the lowest, but the heterozygosity ratio of non-3*n* MS to 3*n* MS in CDS was the highest (Fig 3F, S4 Table). A previous study showed that subdivided populations show a higher effective population size and lower selection coefficient (26). The Central Asia and Siberia population may be composed of subpopulations, which may affect the selection pressure against MS. These results indicate that MS is evolved by the combination of population history and natural selection.

MS has been used to infer genetic structures because of high genetic diversity (27–30). In a previous study, PCA was conducted using 53,002 MS, but the genetic structures by the MS PCA were less clear than that by SNP PCA (18). In the present study, we used a larger number of MS (253,114 MS with MAF ≥ 1 %) and obtained highly concordant results with SNPs (Fig 4 A,B). Although the overall patterns were similar between MS and SNP (Fig 4, S3 Fig), small differences were observed in African and Oceanian populations (Fig 4 C-F). In Oceanian populations, NAN-Melanesians (NAN; Non-Austronesian) and Bougainville, who belong to Melanesians, were clustered in the SNP PCA but not in the MS PCA (Fig 4 EF). In the African populations, Biaka and Mbuti populations showed different patterns between the SNP and MS PCAs (Fig 4 CD), which may be caused by hidden population structures. Although the efficiency of using MS for genetic structures should be evaluated by a larger number of samples, these results indicate that MS can be an additional marker set and may detect hidden population structures in the human population.

In addition to the modern human samples, we analyzed a deep sequenced ancient human sample (F23) (20). In ancient genome sequencing, the DNA fragmentation and library construction process should affect the quality of the sequence reads. To evaluate the quality of the MS call, we compared the distribution of the VAF of this sample with that of modern human samples (Fig 5 A-D). The clear skew of the VAF was observed in MS with unit lengths ≤ 2 bp, suggesting that MS with a short unit are strongly affected by the quality of DNA samples.

However, the distributions of unit lengths ≥ 3 bp were not different, and therefore we used these MS for the analysis. In the PCA, F23 was close to East Asians, which is consistent with the SNP PCA in a previous study (Kanzawa-Kiriyama et al., 2019) (Fig 5 E,F). This result suggests the applicability of MS to ancient human samples.

We found 22 novel highly polymorphic MS for the personal identification. Using the allele frequencies in a Japanese population, the discriminative power was estimated to be 1×10^−17^, which is sufficient for personal identification. Although the discriminative power of our MS set is slightly lower than that of the Globalfiler kit, which is a standard MS set, for a Japanese population (5.6×10^18^) (31), the length of our MS was shorter and can be genotyped by short-read sequencers. Additionally, the PCR success rate of MS is known to be affected by the length of the MS (32), and our shorter MS may be robust to DNA degradation.

This study provides a comprehensive catalog of MS in human populations and shows the applicability of MS to modern and ancient human population studies. Nevertheless, our study has several limitations. First, the genotyping of MS needs reads that cover MS regions. Therefore, the amount of data and read length strongly affect the results. For example, we removed 824,459 MS and 395 samples from the SGDP and HGDP due to insufficient depth. Deeper sequence data would improve the quality of the MS calling. Second, long MS cannot be analyzed using short-read data. A recent study using a long-read sequencer reported high genetic variation in long repeat regions (33). In the future, the application of our algorithm to long-read data should detect a larger number of polymorphic MS.

To conclude, here we analyzed MS polymorphisms using large publicly available human genome sequencing datasets. This study revealed a pattern of MS polymorphisms and identified polymorphic MS in the human population. The comparison of the heterozygosity among populations suggests that MS have evolved by random genetic drift and negative selection. PCA suggests that MS detect the genetic structures of human populations. Currently, large-scale sequencing projects are ongoing worldwide, in which the analysis of MS, in addition to SNPs, should provide deeper understanding of human genetic variations and benefit genome medicine.

## Materials and Methods

### Data

We downloaded the following publicly available sequencing datasets: the Korean Personal Genome Project (KPGP), Simons Genome Diversity Project (SGDP)(18), and Human Genome Diversity Project (HGDP)(19), Japanese samples from the International Cancer Genome Consortium (ICGC)(34), and an ancient DNA sample (20) (S6 Table, S7 Table). The SGDP (n = 300) and HGDP (n = 1064) samples were collected from various populations throughout the world. The KPGP sequenced 107 Koreans. The ICGC performed WGS of cancer and matched normal samples; in this study, we used the WGS data of normal Japanese samples (n = 255). A deep sequenced ancient genome dataset (F23) was also analyzed (20).

KPGP data were used for the MS selection and parameter optimization of the MS calling. The KPGP has data from monozygotic twins (KPGP-00088 and KPGP-00089), which were used for the parameter optimization of the MS calling (S6 Table). One of the monozygotic twins and other Korean samples (in total n = 81) were used for the MS selection. The HGDP and SGDP samples were merged and used as a single dataset (S6 Table). When population names were inconsistent between the SGDP and HGDP, we adopted the population names of the SGDP (S6 Table).

As a result of the sample selection (see below), we selected 81 Korean samples from the KPGP and 969 samples from the HGDP and SGDP (138 samples from Africa, 70 from America, 48 from Central Asia and Siberia, 193 from East Asia, 38 from Oceania, 195 from South Asia, and 287 from West Eurasia). The quality of the ICGC sequencing data was not constant among samples; therefore, Japanese samples from the ICGC were used to estimate the allele frequencies of our MS marker set for personal identification (S7 Table).

The downloaded bam files were results of the mapping to the GRCh37; therefore, our analysis was based on the GRCh37.

### MS genotyping using MIVcall method

Target MS regions were selected in our previous study (13) using three software packages: MSDetector, Tandem Repeat Finder, and MISA software (35–37). Regions were filtered based on the uniqueness of the flanking sequences and the distance to other MS. Insertions and deletions in a target MS were detected using the MIVcall method (13); MIVcall counts the length of each MS in each read. When multiple lengths are observed in a MS locus in a sample, the most frequent pattern is assumed to be present, and the second most frequent pattern is examined. The likelihood value was calculated based on the number of reads. Genotypes are determined based on the likelihood value, the number of reads that support the pattern, and the VAF.

### Establishing the MS detection method

In our previous study (13), the optimal criteria of likelihood and number of reads (the minimum - log10(likelihood) value and minimum number of reads for allele detection) were chosen for analyzing somatic mutations. To obtain the optimal criteria for a polymorphism, we used monozygotic twins in the KPGP (KPGP-00088 and KPGP-00089). Since all genotypes of monozygotic twins are identical, we tested various parameter sets and compared the concordance rates of genotypes between twins.

### Sample selection and MS filtering

Since MS are susceptible to sequencing errors, selecting high-quality samples is necessary. Thus, MS covered by less than 10 reads were considered MS with insufficient depth. We excluded samples if more than 4% of MS loci had insufficient depth.

Next, we selected MS loci from the 9,292,677 MS selected in our previous study (13). Using the 81 Korean samples in the KPGP, we counted the number of samples with insufficient depth for each MS and removed samples if the percentage of MS with insufficient depth was ≥ 15%. Additionally, we tested deviations from the HWE with Fisher’s exact test for 2 x *n* contingency table (*n*; number of genotypes) in the KPGP (*α* = 0.0001).

### Genome-wide pattern of MS

To reveal the landscape of MS polymorphisms, we analyzed the features of MS. This analysis was performed using HGDP and SGDP samples. We analyzed the association of the length of a MS region in the reference genome with the number of samples with insufficient depth and heterozygosity. We also examined the number and heterozygosity of MS for repeat units with different sequences (for example, A, G, and AC). In this analysis, we merged MS with different unit sequences if reverse-complement (e.g., GT to CA) or reverse (e.g., TA to AT) generated the same sequences.

We then focused on MS in CDS and in non-CDS. We compared GC content, lengths of MS, number of alleles, and heterozygosity between CDS and non-CDS MS. In CDS, MS were classified into 3*n* MS (3 and 6 bp) and non-3*n* MS (1, 2, 4, 5 and 7 bp). We compared heterozygosity among all MS, 3*n* MS, non-3*n* MS, MS in CDS, and MS in non-CDS. The ratios of the mean heterozygosity of non-3*n* MS to the mean heterozygosity of 3*n* MS were compared among the different populations. We also performed a pathway analysis for genes with multi-allelic non-3*n* MS using the Reactome database.

### Analysis of population structure with MS

We conducted five dimensionality reduction methods: Principal Component Analysis (PCA), t-Distributed Stochastic Neighbor Embedding (t-SNE), Uniform Manifold Approximation and Projection (UMAP), PCA-t-SNE, and PCA-UMAP. For these analyses, the genotypes of MS and SNPs were converted to numerical values. SNP genotypes were converted to numerical values by counting the number of minor alleles. For MS, we used two methods, the multiallelic method and average method. In the multiallelic method, each allele at a MS locus is treated as a different marker (if three samples have the following genotypes, [8/8, 13/14, 8/14], we converted them into 3 independent pseudo-loci: [2, 0, 1] (8 or else), [0, 1, 0] (13 or else), and [0, 1, 1] (14 or else)) (38). In the average method, we calculated the average length of two alleles in each individual (in the previous example, the average method converts the genotypes to [8, 13.5, 11]) (S5 Fig). We performed a PCA with both methods and compared the results.

We used the MS and SNP with a MAF ≥ 1% in the SGDP and HGDP samples. MAF was calculated by 1 – (major allele frequency). A PCA was conducted for all samples and samples in each region (Africa, America, Central Asia and Siberia, East Asia, Oceania, South Asia, and West Eurasia). Other dimensionality reduction methods (t-SNE, UMAP, PCA-t-SNE, and PCA-UMAP) were applied to all samples only.

### MS set for personal identification

Individual identification with MS is an important topic in human genetics. We selected a set of 22 MS and estimated the discriminative power. We calculated the heterozygosity of MS with repeat unit lengths = 4 and selected 22 MS with the highest heterozygosity in each chromosome. The allele frequencies of the selected 22 loci were estimated using 255 Japanese samples from the ICGC data. The discriminative power was calculated as the product of frequencies of the most frequent genotype in each locus (31).

### Analysis of ancient DNA samples

To analyze MS variation in ancient DNA, we analyzed one ancient DNA sample with a higher depth of coverage from a previous study (Sample ID; F23) (20). To examine the quality of the variant calling, we calculated the VAF of each MS of F23 and compared it with the average VAF of 10 randomly selected HGDP samples. We then conducted a PCA for F23 sample with the SGDP and HGDP samples.

### Programming languages

We used Python (https://www.python.org) for this study. PCA, TSNE, UMAP, and Decision Tree were conducted with the sklearn package.

## Acknowledgements

This work was supported by Grant-in-Aid for Scientific Research on Innovative Areas from JSPS grants (Grant Number 18H02680 to A.F.), Yaponesian genome MEXT KAKENHI (Grant Number 18H05511, to A.F.), and the AMED (Grant Number JP21km0908001, to A.F.). Data analysis was performed using the SHIROKANE supercomputer (https://supcom.hgc.jp) at the University of Tokyo and the NIG supercomputer at the National Institute of Genetics.

## Author Contributions

Study design: A.F. Data analysis: L.G. and A. F. Manuscript writing: L.G. and A. F. Interpretation of data: L.G., Y. K. and A. F.

## URLs

KPGP sequencing data: http://kpgp.kr.

The Reactome database: https://reactome.org/PathwayBrowser/

