## Supplementary material for "Comprehensive analysis of microsatellite polymorphisms in human populations": S_Figs

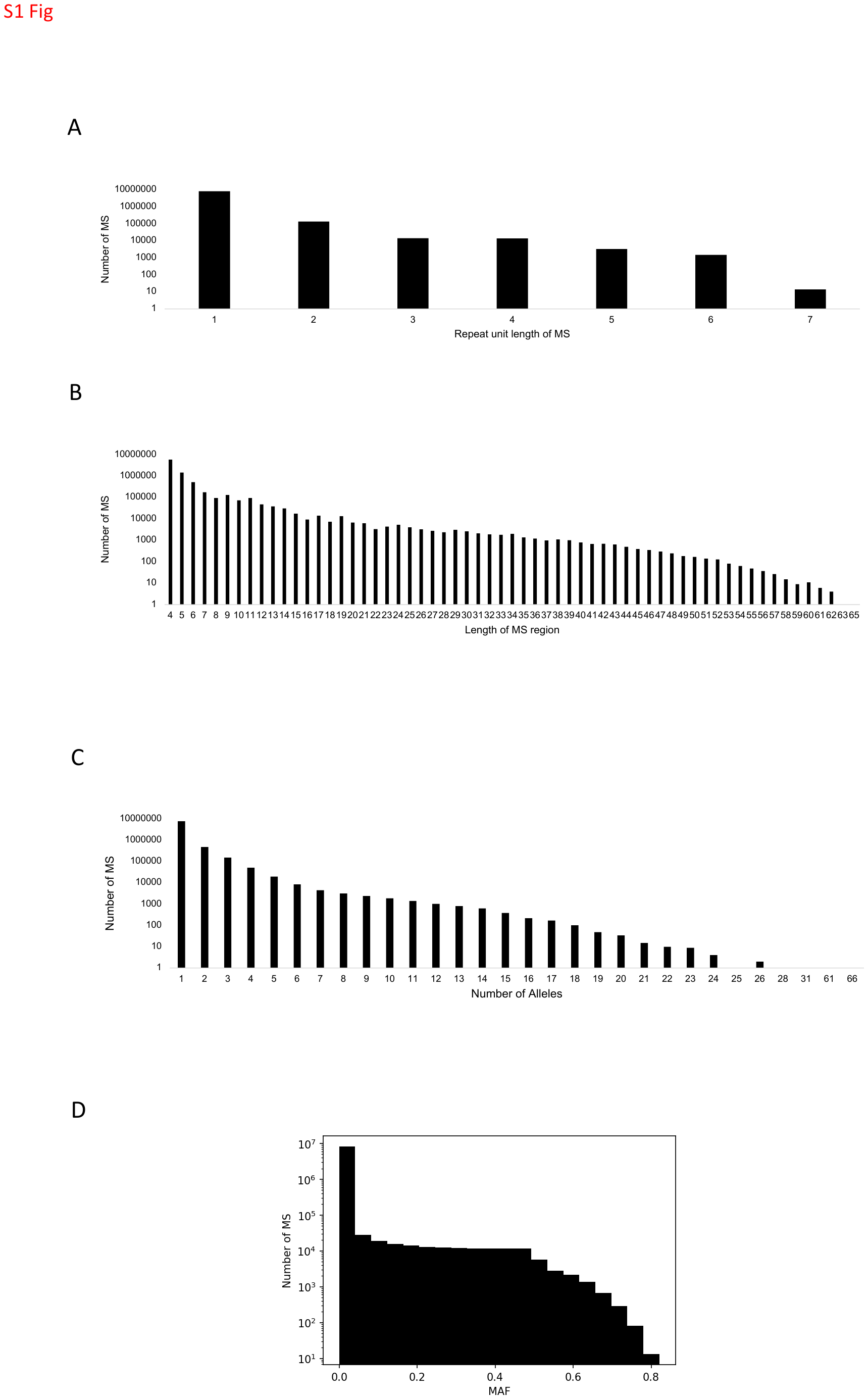

S1 Fig1.

Distributions of MS. (A) Distribution of repeat unit lengths. (B): Distribution of lengths of MS regions in the reference genome. (C): Distribution of number of alleles. (D): Distribution of minor allele frequencies (MAF).

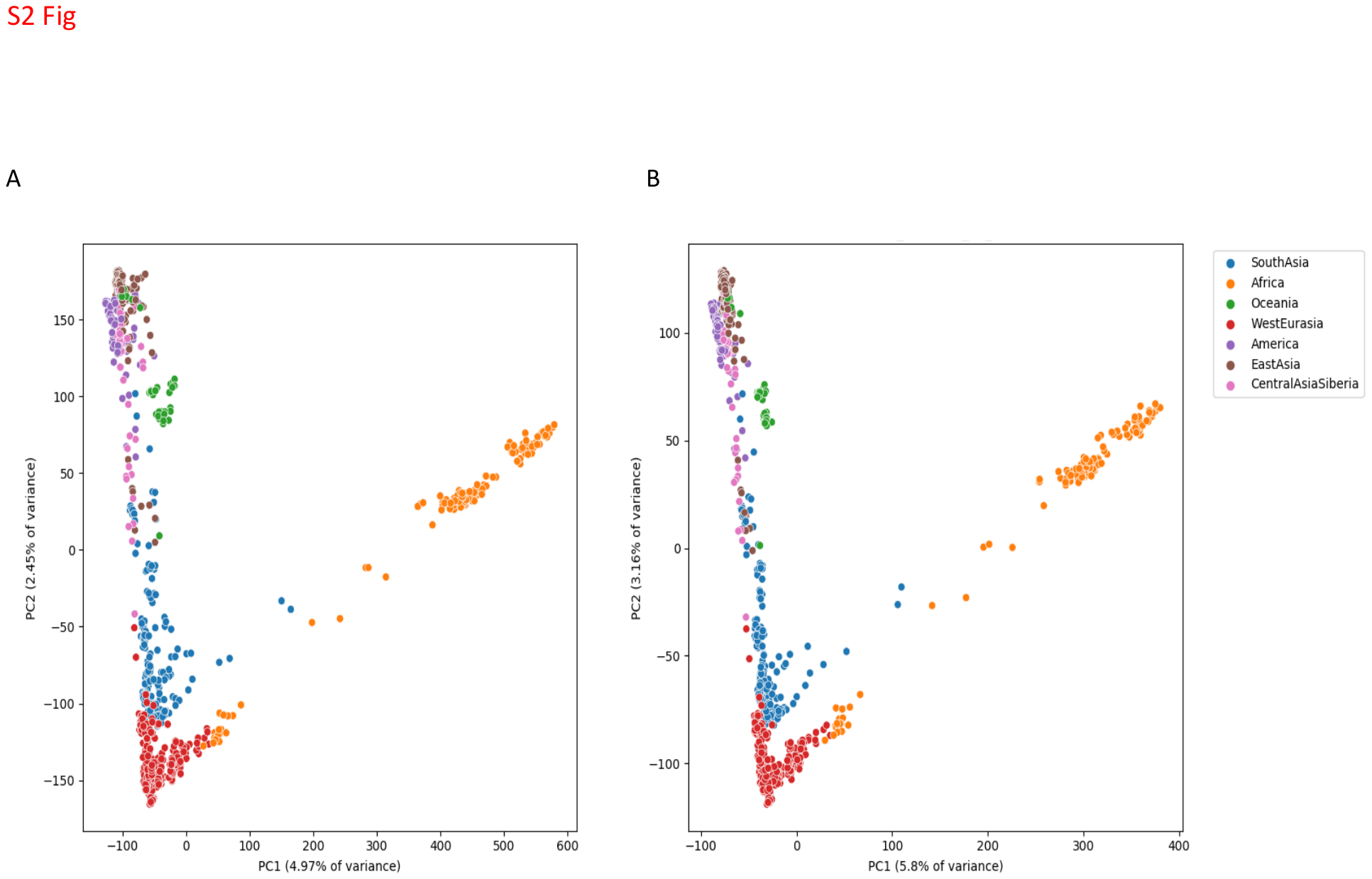

S2 Fig.

PCA with SGDP+HGDP samples using two methods. (A) Multiallelic method. (B) Average method.

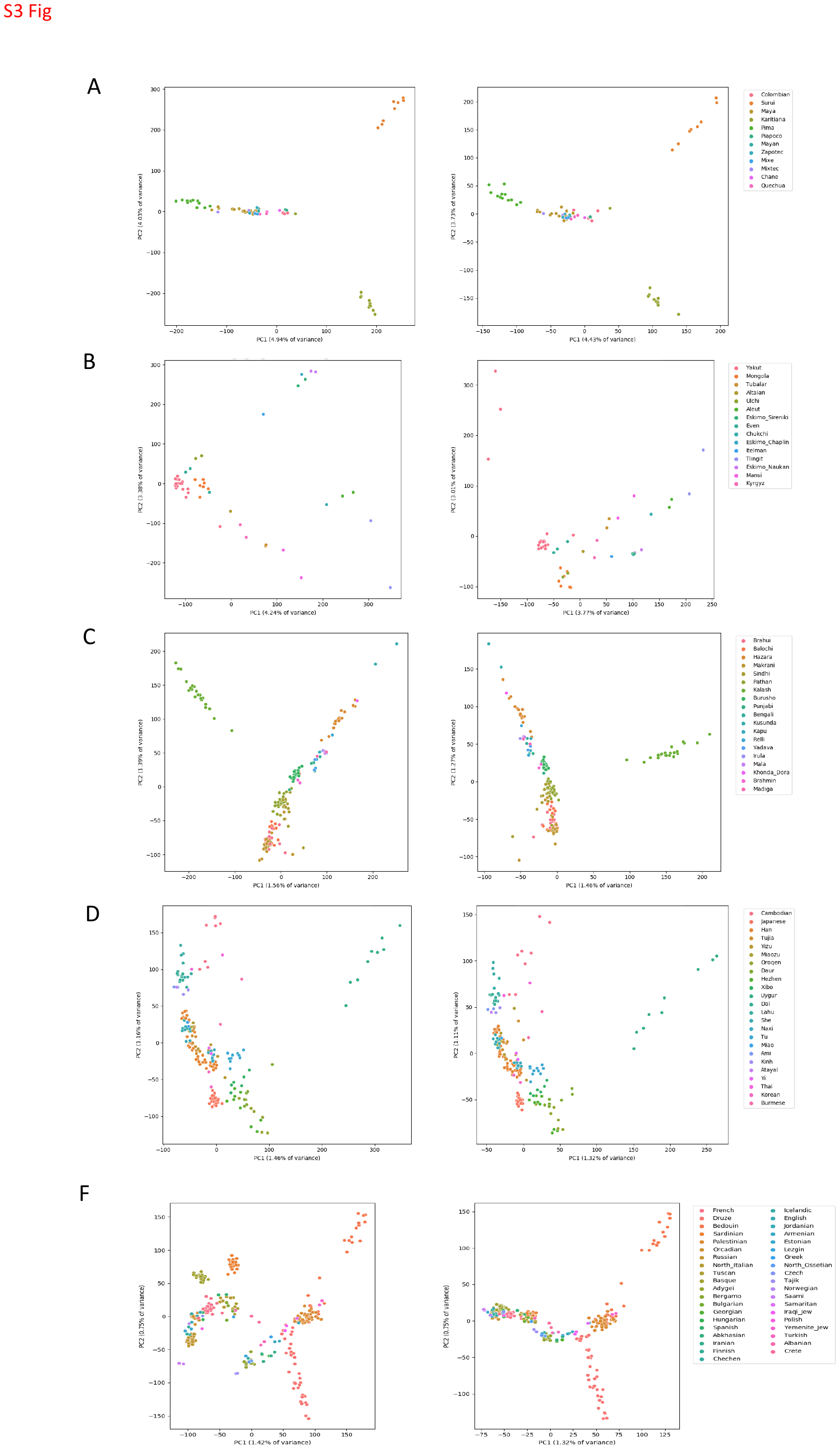

S3 Fig.

PCA of all SGDP and HGDP samples using SNP (left) and MS (right). (A) America. (B) Central Asia and Siberia. (C) South Asia. (D) East Asia. (E) Oceania. (F) West Eurasia.

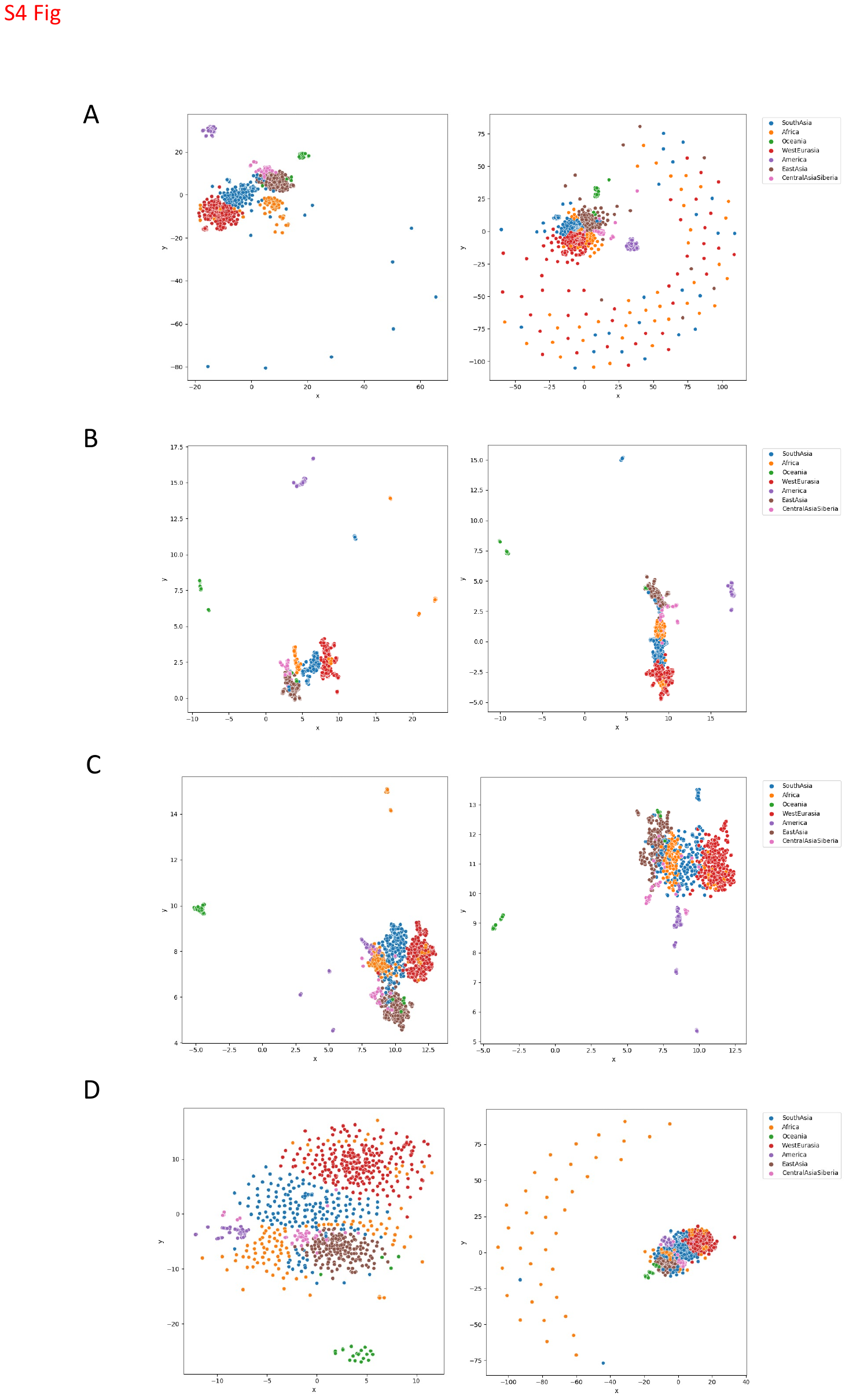

S4 Fig.

Dimensionality reduction of all SGPP and HGDP samples using SNP (left) and MS (right). (A) t-SNE. (B) UMAP. (C) PCA-t-SNE. (D) PCA-UMAP.

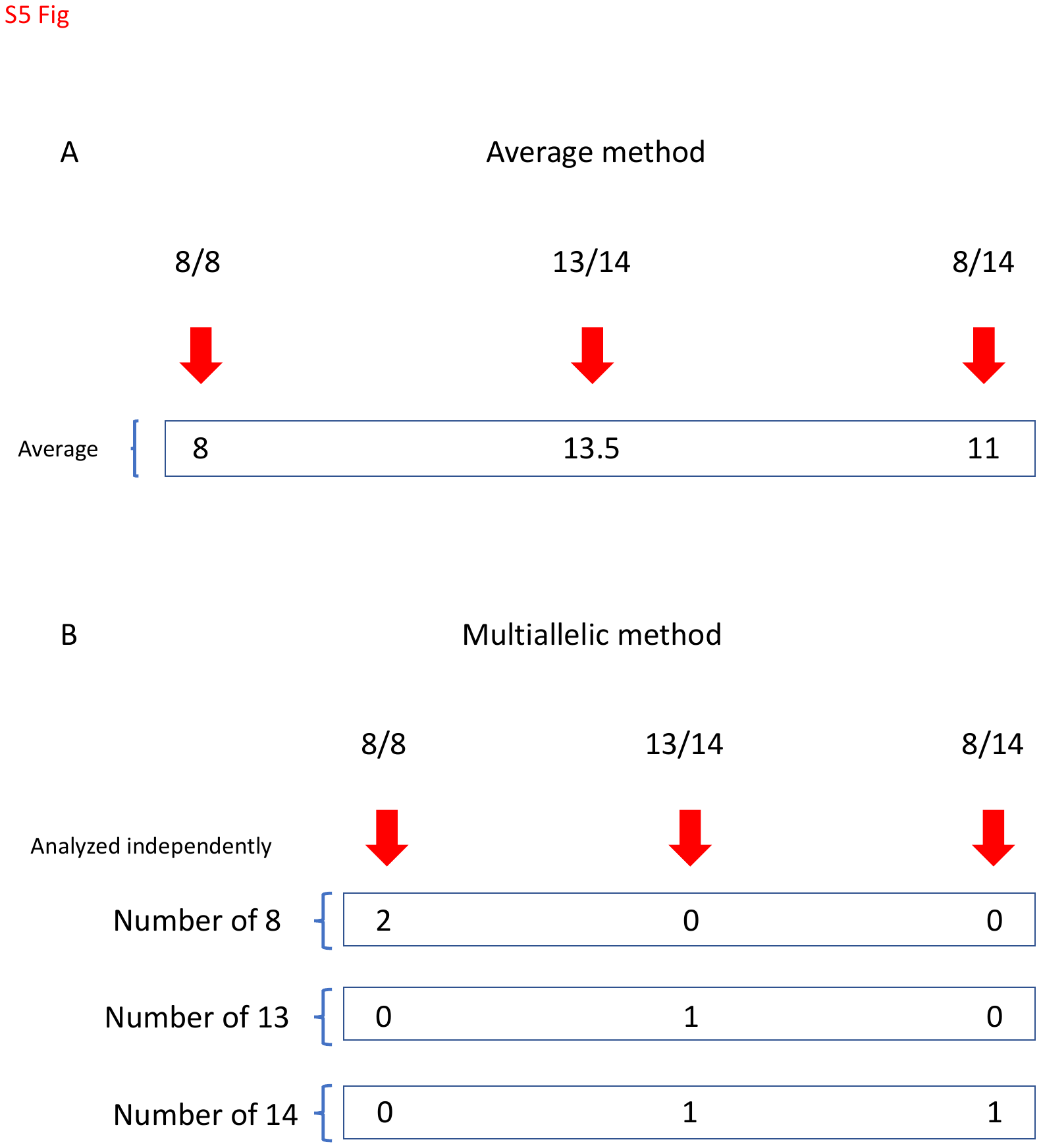

S5 Fig

Methods to convert MS genotypes to numerical values.

(A) Average method. (B) Multiallelic method
